# Resolving the connectome – Spectrally-specific functional connectivity networks and their distinct contributions to behaviour

**DOI:** 10.1101/700278

**Authors:** Robert Becker, Alexis Hervais-Adelman

## Abstract

The resting brain exhibits spontaneous patterns of activity that reflect features of the underlying neural substrate. Examination of inter-areal coupling of resting state oscillatory activity has revealed that the brain’s resting activity is composed of functional networks, whose topographies differ depending upon oscillatory frequency, suggesting a role for carrier frequency as a means of creating multiplexed, or functionally segregated, communication channels between brain areas. Using canonical correlation analysis, we examined spectrally resolved resting-state connectivity patterns derived from MEG recordings to determine the relationship between connectivity intrinsic to different frequency channels and a battery of over a hundred behavioural and demographic indicators, in a group of 89 young healthy participants. We demonstrate that each of the classical frequency bands in the range 1-40Hz (delta, theta, alpha, beta and gamma) delineates a subnetwork that is behaviourally relevant, spatially distinct, and whose expression is either negatively or positively predictive of individual traits, with the strongest link in the alpha band being negative and networks oscillating at different frequencies, such as theta, beta and gamma carrying positive function.

## 1. Introduction

Even in the absence of any task, the brain expresses patterns of functional connectivity. Such resting-state activity is thought to be determined by intrinsic features of the underlying neural architecture^1^. Thus, spontaneous activity of the brain provides an insight into endogenous features, which are increasingly understood to be substantial contributors to individual differences in behaviour. A number of recent studies have highlighted the extent to which resting state connectivity metrics reveal individual “wiring patterns” of the brain that are significantly predictive of behavioural, and even demographic features. Notably, resting state fMRI (RS-fMRI) connectivity patterns have been shown to be predictive of task-related activation patterns^2^ and even to constitute uniquely identifying fingerprints that can distinguish between individuals^3^. Such findings suggest that resting-state activity patterns reflect a neurobiological framework that determines a large degree of individual differences in brain responses to tasks, and of individual differences in behaviour. In a sense, they may be considered the brain’s “priors”^4^ for determining how it responds to input.

A recent effort to relate whole-brain resting state fMRI connectivity to individual differences in behaviour employed canonical correlation analysis (CCA) to establish how inter-individual differences in connectivity relate to differences in a broad battery of behavioural and cognitive variables^5^ (non-imaging subject measures, behavioural measures), using a sample of 461 participants from the Human Connectome Project^6^. CCA was used to find the maximal correlations between combinations of variables in the connectivity and subject measures, revealing a canonical mode of covariation that is highly positively correlated with positive cognitive traits as well as desirable lifestyle indicators. This is a fascinating finding that highlights the extent to which large-scale neural connectivity of the human brain is functionally relevant. However, this discovery invites further and deeper inquiry: fMRI is rightly prized for its high spatial resolution, but its temporal resolution is inherently limited by the sluggishness of the haemodynamic response that it records. Consequently, fMRI cannot be used to discriminate between neural processes that occur at timescales in the sub-second range. These are, however, of profound relevance to further elucidating the nature of resting state brain activity, and its relationship to behaviour.

A characteristic feature of the human brain’s electrical activity at rest is the presence of oscillatory signals. Oscillations are the product of repetitive or cyclical patterns of brain activity, which are observed to occur at different frequencies. Oscillations exist at multiple scales, spatially and temporally. Spatially they range from single neuron oscillations, to local field potentials and large-scale magneto-electric brain signals. Temporally, different oscillatory frequencies are thought to represent different channels in which neural activity is communicated, and might therefore fulfil different functional roles^7^. Even at a relatively coarse level of spatial resolution, an extensive literature of M/EEG or electrophysiological studies has established how neuronal processes occurring in different frequency ranges have different functional associations - from the relatively slow sleep rhythms (e.g. delta 1-3Hz waves in NREM sleep^8^) to memory processes in the hippocampus^9^ (mostly in the 4-7 Hz theta range), to processes related to sensation (covering the alpha and beta range of 8-30 Hz^10,11^) or executive functions (e.g. working memory^12-14^) and feature binding during visual and other task processing^15,16^, as reflected in higher frequencies such as the gamma frequency range, i.e. 30Hz and above.

Oscillations can be characterised in terms of frequency, amplitude and phase. Much of the research into the functional relevance of neural oscillations has focused on phase or amplitude, which have been shown to index different states of neuronal excitability^17-23^ and ultimately, to predict behavioural and perceptual performance in a variety of experimental task conditions, domains and sensory systems^24-27^. While such research has been fruitful, if we are to consider different oscillatory frequencies as carriers of different information, an analysis of the networks that they constitute is necessary.

Synchronised patterns of modulation of activation at separate loci are believed to reflect coupled activation, and consequently to reveal functionally connected areas, and ultimately, functional networks. By examining resting state activity in distinct frequency ranges, it is possible to define large-scale brain connectivity networks that are frequency-resolved. Recent efforts have identified large-scale (i.e. whole-brain) electrophysiological networks with topographical and spatial structures comparable to the resting state networks established by fMRI studies. For example, the motor networks or visual components, analogous to those previously described in fMRI^28,29^ have been reported in MEG^30-32^. Interestingly, these networks, while spatially similar to fMRI-derived RSNs, had distinct spectral properties. That is - they exist as coherent activity in different frequency regimes: the motor network being most pronounced in the beta range (14-30 Hz) whereas networks attributable to visual areas were most visible in the alpha range 8-12 Hz. The finding of different topographies in spectrally distinguished networks hints at the potentially different function these spontaneous networks might serve, but beyond the more intuitively interpretable sensory- and motor-related networks mentioned above, the roles and particular functional relevance of the spatial networks that emerge at different frequencies, remains elusive. At this point, it is worth highlighting that all neural communication is constrained by the underlying anatomical connectivity, and that any network of information transfer that is discovered is necessarily a reflection of intrinsic brain architecture. Consequently, any relationship elucidated between functional connectivity patterns and behaviour depends upon neuroanatomy, but they are not fully determined by it^33^ – the existence of connectivity in different frequency ranges is highly suggestive of the possibility that the anatomically constrained connections can be exploited to multiple ends.

The precise role of network communication at different frequencies is the subject of ongoing and increasing interest. One overarching notion is that of “multiplexing”, *viz*. that different oscillatory frequencies comprise different communication channels allowing the same anatomical connections to be used for different functional roles^34^. An influential theory with a very broad scope - ‘communication through coherence’ (CTC) - proposes that the observed frequency-constrained networks, i.e. networks that share a common rhythm on a carrier frequency, reflect activity of cell assemblies that are in communication with each other – e.g. when binding different sensory modalities together^15^. In particular, coherence in the faster gamma band (40 Hz and above) is well examined and seems to confirm the proposals of CTC theory. CTC also affords the existence of similar mechanisms in other frequency bands, although these are less well explored and less is known about their likely functional role or roles.

Another recent focus of enquiry, underscoring the importance of neural activity in different frequency bands has been how different frequencies reflect the activity of neurons having different roles in the directionality of neuronal communication. For example, it is known that bottom-up processes are preferentially mediated by higher-frequency (e.g. gamma) activity, while top-down processes have principally been associated with rhythms in lower frequencies^35^. In sum, there is compelling evidence indicating that spectrally confined, oscillatory activity may serve different purposes in network communication, be it in directing information flow or in circumscribing the neural assemblies that preferentially operate as coherent ensembles.

Despite these advances, a holistic analysis of the functional role of oscillatory networks, taking into account the full spectral richness of the underlying constituents of human spontaneous brain activity, and its relationship to an extensive range of behavioural factors has, so far, been beyond our reach. Here, we seek to expand upon and complement the existing rs-fMRI findings by exploring the relationships between the same set of Subject measures and spectrally-resolved connectomes derived from rs-*MEG*, also made available as part of the HCP dataset^36^.

Specifically, we address the following questions. Firstly, can we identify and characterize a global mode linking brain connectivity and behaviour using spectrally resolved MEG? Secondly, is this mode comparable to that revealed for rs-fMRI? And thirdly, what – if any - additional information can we glean from the spectral richness of MEG afforded by its high temporal resolution? By exploring these questions, we will shed light upon the spatial structure and functional contributions of the spectral components of resting state connectivity and how they relate to a broad spectrum of behavioural indicators.

## 2. Results

### 2.1. MEG-derived functional connectivity reveals a global brain-behaviour mode along a positive-negative axis of subject measures

We used canonical correlation analysis (CCA) to examine the relationship between spectrally resolved resting-state MEG connectivity and a battery of 131 behavioural, demographic and personality variables in 89 healthy young individuals. CCA is a method that identifies an optimal linear combination of features (modes) in two separate data sets that maximizes correlation between them – in our case functional, spectrally resolved MEG connectivity and subject measures. MEG connectivity was collapsed into five conventional frequency bands in the range 1-40Hz (delta [0.5 - 3 Hz], theta [3-7 Hz], alpha [7-13 Hz], beta [13-25 Hz] and lower gamma [25-40 Hz]). Connectivity was determined for the whole brain divided into 100 functionally defined parcels (see Supplementary Figure 1 and Supplementary Table 1). To remove simple effects of resting state power, average resting state spectral power was regressed out of connectivity strength variations across subjects in a pairwise manner, i.e. for each connection, and frequency band, power of both nodes was regressed out. CCA was then performed on a PCA reduced subspace of this data, examining the potential link between subject-wise variations of connectivity strength and behavioural performance (see Methods), identifying one significant canonical mode after permutation testing (r = .94, p <= 10^−4^, see also Figure 1S4 for a visualization of the CCA approach). This mode links spectrally resolved brain connectivity and behaviour. The existence of a significant mode indicates that there is a significant relationship between resting state connectivity and the subject measures.

**Figure 1.**
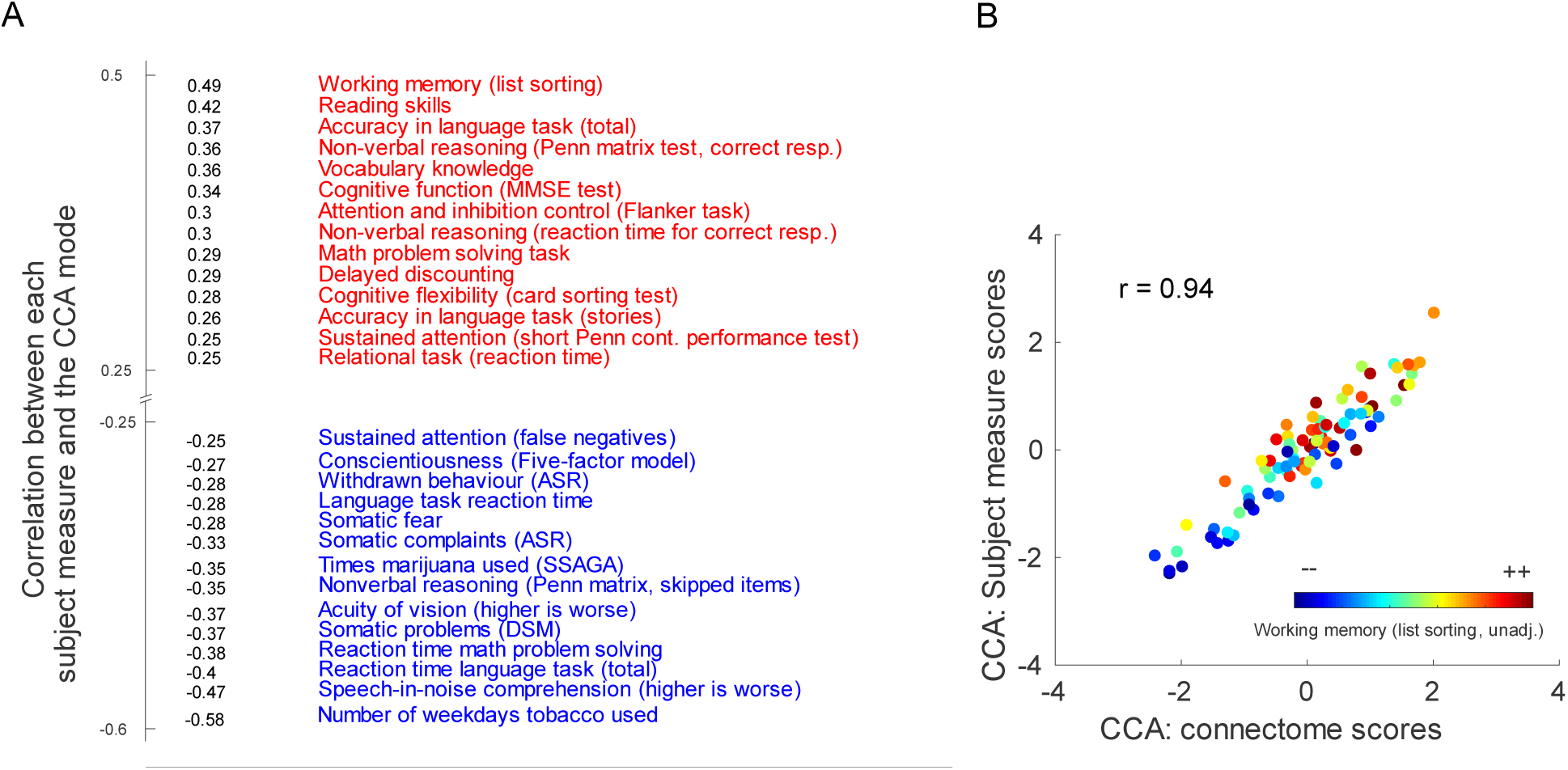
Canonical correlation analysis of spectrally resolved MEG data and a large set of behavioural and other subject measures results in one significant mode (r = 0.94, p < 10^−4^, corrected for multiple comparisons by permutation testing). **A**. The canonical mode arranges behavioural variables and subject measures on a positive-negative axis, similar to what has been previously reported for hemodynamic measures of brain connectivity^5^. At maximum positive correlation there are mostly subject measures indexing cognitive performance such as reading skills and vocabulary knowledge, while on the negative end of the spectrum are subject measures like somatic problems, and tobacco consumption (thresholded at a correlation coefficient of |r|>.25). **B**. Correlation of CCA-derived subject measure scores and connectivity scores for the first canonical variates identified, i.e. the first canonical mode. In color the behavioral score for the working memory test is shown per subject. Abbreviations – Penn matr. = Penn matrix test; ASR: Achenbach Adult Self Report; DSM: Diagnostic and Statistical Manual, MMSE = mini mental state exam, SSAGA = Semi-Structured Assessment for the Genetics of Alcoholism, unadj = unadjusted for age effects. Please note that some secondary measures (e.g. similar metrics for tobacco consumption are left out to avoid redundancy).

In order to provide a functional characterization of this mode of maximum covariation between subject measures and connectivity, we evaluated how the mode (i.e. the observed first canonical variates) covaries with the individual subjects’ scores on each of the subject measures and connectivity measures. This procedure reveals the way in which the mode that best aligns the full set of subject measures and the connectivity patterns explains the inter-individual variation in each of the subject measures. Consequently, positive correlation between an SM and the canonical variate suggests that the higher a subject ranks on this subject measure, the larger this subject’s score for the canonical variate. Conversely, a negative relationship between an SM and the canonical variate indicates that higher scores on that SM are associated with lower subject scores on the canonical variate.

The behavioural variables most strongly positively and negatively associated with this mode are shown in Figure 1A. The subject measures are arranged along a positive-negative axis that is qualitatively the same as that previously described for an fMRI derived CCA mode^5^. It ranges from subject measures that entail tobacco consumption and somatic problems (which may quite reasonably be characterised as ‘negative’) to subject measures indicating high cognitive performance (at the other end of this ranking, interpreted as ‘positive’). The nature of the positive relationship between the mode and performance on a working memory task is further illustrated in Figure 1B, where a clear pattern emerges demonstrating that higher individual scores on the subject measure and connectome canonical variate are predictive of superior list sorting performance.

This outcome alone is remarkable – despite using a different imaging modality and having access to only a substantially smaller sample (89 vs. 461 data sets) the nature of the mode revealed is highly consistent with that previously demonstrated for the same set of subject measures and rs-fMRI connectivity. A quantitative comparison with rs-fMRI based CCA results, as well as the impact of parcellation scheme and subdivision into the selected frequency bands on the outcome is presented in Figure 1 S5.

Although this MEG-derived canonical mode resembles and partially even reproduces a previously identified fMRI-derived brain-behaviour mode, we are at pains to point out that this does not automatically render it *the* ‘universal’ brain-behaviour mode. Given a different array of subject metrics, or a qualitatively different array of brain metrics, the principal brain-behaviour mode uncovered by CCA could be different to the one at hand. This possibility is, however, speculative. What we may state with confidence is that, independent of the underlying imaging modality, a canonical brain-behaviour mode, optimally matching spectrally resolved connectomes and subject measure, can be identified, which arranges subjects on a positive-negative axis consistent with a study in which connectivity was derived from hemodynamic measures of brain activity.

### 2.2 The CCA Mode is Composed of Spatio-Spectrally Segregated Subcomponents

In a second step, we characterize the connectivity components of the identified brain-behaviour mode, for each of the five frequency bands and their connectomes. In order to do this, the connectivity patterns across subjects in each band were correlated with the individual connectome weights of the significant CCA mode. This reveals which connections (hereafter referred to as edges) of the connectome are more, or less, related to the mode, and therefore associated to individual differences over subject measures. The outcome of this analysis is shown in Figure 2. The correlation coefficients calculated in this analysis will be referred to as edge modulations (EMs). The patterns of within-band edge modulations (EMs) significantly related to the mode are clearly differently distributed over the brain and have distinctly negative or positive relationships with the mode in each of the analysed frequency bands.

**Figure 2.**
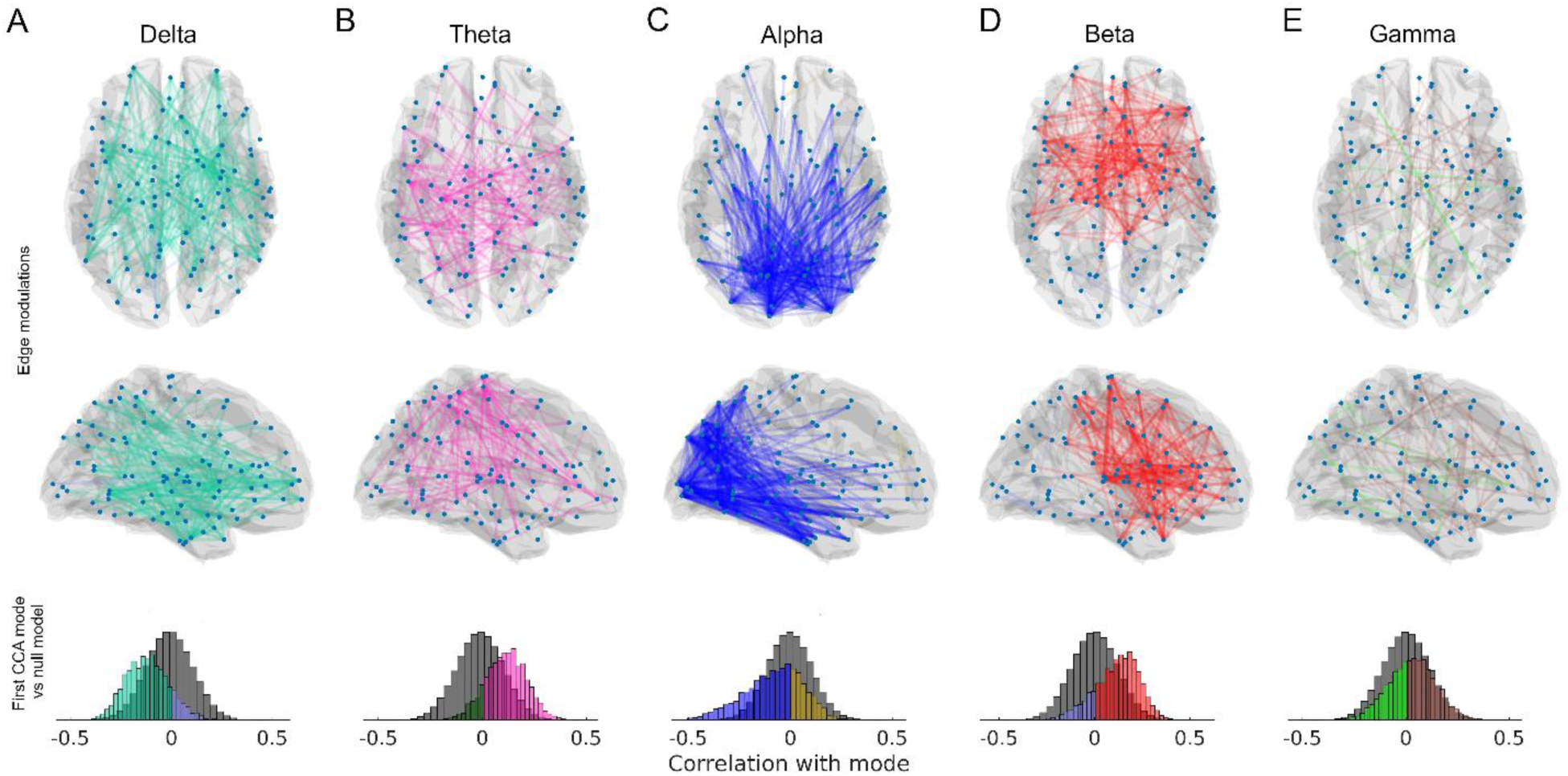
**A-E**. Edge modulations (i.e. edges whose connectivity on a subject-by-subject level covaries with behaviour) for each frequency band, from delta, theta, alpha and beta to gamma band. Edge modulations of frequency bands are colour coded (2 colours per band, positive and negative, encoding the correlation coefficients, in a range from -.5 to .5 to the observed canonical mode). Edges rendered on the brain templates are thresholded at the 0.5^th^ bottom and 99.5^th^ top percentile (of the permutation-based null distribution) for visualization. Each of the bands shows a preference for either positive or negative relationship to the mode, but not a mixture of both. This is also visible in the histograms in the bottom row that depict the distributions of all (i.e. unthresholded) correlation coefficients, with comparison to null distributions generated by a permutation test (n=10000, in grey).

Figure 3A shows the top 50 edge modulations (EMs) across all five frequency bands, providing an overview of the diversity of connectivity patterns in each of the frequency ranges analysed. To reveal the relative functional importance of each node in terms of its relationship to the mode, averaging was performed over all the EMs in which the respective node is involved. We thus derive maps of accumulated EMs, which are shown in Figure 3B. As described above, the distributions of EMs across bands is primarily unipolar, each of the five frequency bands preferentially showing either a positive or a negative relationship with the mode, but not both (Figure 3B).

**Figure 3.**
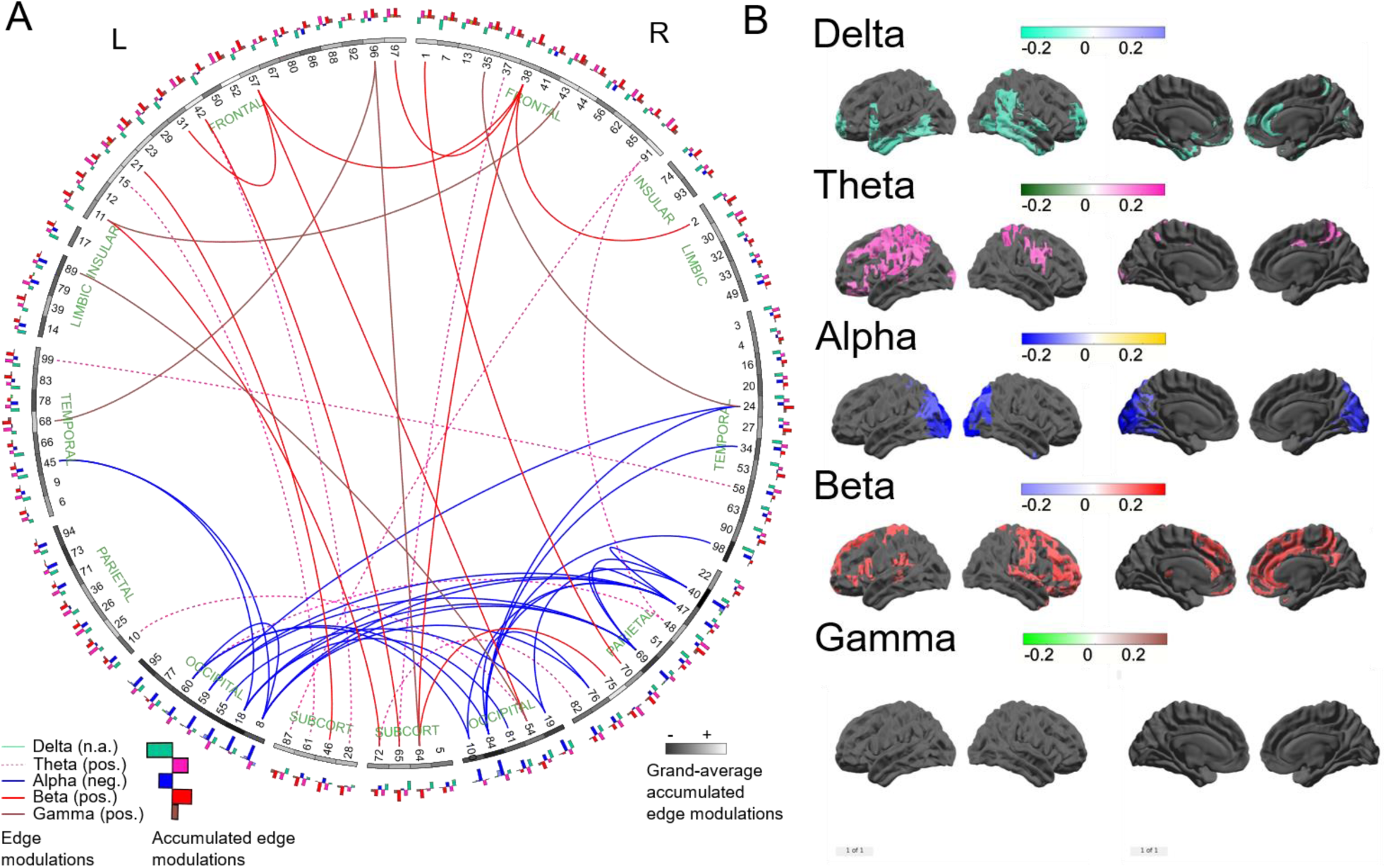
**A**. Connectogram-style visualization^37^ of the EMs presented in Figure 2. This figure shows the top 50 EMs (i.e. the top 50 positive and negative edges, respectively), across all five frequency bands (globally tresholded across all bands pooled). Nodes are represented on the outside of the ring, in an approximately anatomically-faithful anterior-posterior and left-right arrangement. Anatomically labeled parcel identities are listed in Supplementary Table 1. Each parcel’s grayscale value indicates its average relationship to the CCA mode, i.e. its average accumulated edge modulation across frequency bands – white indicates a positive relationship to the mode, and black indicates a negative relationship (averaged over bands and all edges, i.e. connections of that parcel). The 5-band histograms on the outer ring indicate the band-specific accumulated edge modulations for each node (same color coding as for the top edges in 1A and the maps in 1B, histograms pointing outwards are positive, inward histograms indicate negative accumulated EMs). **B**. Maps of accumulated EMs for all frequency bands. These represent the average EM value, over all connections, for each parcel. Thus, these accumulated maps represent the overall involvement of each parcel in predicting behaviour and provide complementary information to the EMs, by showing the presence of connectivity foci, i.e. nodes of particular importance to the EM network, acting as sinks or hubs. Thresholded at the 10^th^ and 90^th^ percentiles for negative and positive accumulated EMs, respectively. NB: No supra-threshold accumulated EMs are observed in the gamma band. Abbreviations: L = left, R = right, pos. = positive, neg. = negative, SUBCORT = subcortical.

### 2.3 Connectivity of spectrally-resolved brain activity is preferentially ‘good’ or ‘bad’, but not mixed

The discovery of frequency-specific connectivity components that correlate either negatively or positively with the CCA-derived mode invites the conclusion that some connectivity patterns are “good”, while others are “bad”. This interpretation is somewhat consistent with previous reports from rs-fMRI data^5^, which also showed that certain subcomponents (especially areas pertaining to the default mode network) of the whole-brain connectome are positively, and others (mostly visual areas) negatively, associated with the positive-negative axis exposed by CCA.

#### 2.3.1 Delta-band and alpha-band EMs show negative relation to cognitive performance

Alpha and Delta-band connectivity were almost exclusively negatively associated with the CCA mode, i.e. negatively correlated with cognitive performance. The parcels with the highest incidence of significant connectivity-behaviour relationships in the alpha-band are principally concentrated in visual-posterior areas, consistent with the usual distribution of alpha power at rest. Given that spectral power was regressed out of these analyses (see Methods), we may conclude that higher *connectivity* in the alpha band, absent the biasing effects of the typically higher power in this frequency range compared to the rest of the spectrum, is predictive of lower cognitive performance and higher incidence of negatively associated health and behaviour indicators. Delta-band connections also correlate negatively with behaviour. The majority of implicated nodes are differently distributed compared to alpha-band EMs, incorporating bilateral inferior posterior and anterior temporal lobes and inferior and orbitofrontal areas.

The negative relationship of the alpha-band posterior EMs to the canonical mode is comparable to the accumulated EMs known from fMRI derived CCA analysis^5^. By means of spectrally resolved CCA analysis, we can assign a neuro-spectral profile to some of these previously reported EMs. For example, the fMRI-derived negative-posterior EMs in early visual areas^5^ are putative equivalents of the MEG-derived negative-posterior alpha-band EMs, exhibiting similar functional relevance (i.e. higher connectivity = lower score in the CCA mode).

The components residing in the delta-band are less easy to map onto previous results. The finding of delta-band specific edge modulations negatively related to the mode is suggestive of a negative impact of increased low-frequency neural coupling on cognitive function. Usually, pronounced delta oscillations are found during sleep, while in the awake human brain, delta is most often recorded in pathological situations such as in the presence of lesions or tumours. Unusually elevated delta coupling may also be a sign of slowed theta oscillations, a phenomenon often observed in pathological cognitive decline, such as Alzheimer’s disease, caused by cholinergic loss^38^. While we do not propose that the young, healthy cohort examined here suffered from any neurological pathologies, it is intriguing to note the fact that slow oscillatory activity is not typically associated with optimal brain function during wakefulness.

#### 2.3.2 Theta-, beta- and gamma band EMs show positive relationship to cognitive performance

Beta, Theta and Gamma-band EMs exhibit a substantially different relationship to the canonical mode, compared to Alpha and Delta. This is the case both in terms of the polarity of the relationship, and the spatial distribution of the relevant nodes. For the observed canonical mode, subjects scoring high in frontal and prefrontal beta-band connectivity also score highly in the identified canonical mode, indicating higher cognitive performance. The patterns that arise from beta-band accumulated EM maps, especially in medial areas, show good correspondence to previous results for fMRI connectivity^5^. For theta EMs and accumulated maps, spatially we see a more pre-central pattern with a functionally similar interpretation – the more pronounced the connectivity is in these edges, the higher the subject scores in the canonical mode, i.e. the higher the subject is ranked on the positive-negative axis identified by the significant CCA mode. The properties of gamma-band EMs are particularly noteworthy. While the overall gamma EM network is somewhat patchy, featuring what appear to be a greater number of short-range connections, its connectivity is still relevant for the global mode of behaviour. It also seems to have fewer sinks or hubs where connections emerge or terminate, resulting in the absence of significant accumulated EMs, in contrast to the more clustered network patterns in frequency bands at lower frequencies. Considering that gamma in general is known to be an index of local activation, offering often relatively fine spatial resolution (e.g. down to mapping the representation of individual digits of the hand^39^), this is not altogether surprising.

This striking dichotomy of either good or bad connectivity – when looking at spectrally resolved connectomes – is both novel and unexpected. Previous results – using fMRI connectivity that is agnostic to the underlying frequencies of neuronal communication – have so far only shown patterns of mixed positive or negative relationship to the observed brain-behaviour mode. We tested whether this segregation does arise from the CCA trivially by creating EMs from surrogate data, by permuting the subject-labels and recomputing the EMs. The resulting null-distributions of correlation coefficients are shown in Figure 2 (bottom row); they do not exhibit any polarity preference or skewedness as the actual EMs do.

### 2.4 Resolved MEG connectivity model shows results comparable to fMRI connectivity

In terms of the resulting canonical mode and its maximum alignment of connectivity and subject measures, the MEG connectivity model used here (5 bands, 100 parcels) shows comparable results with regard to the results obtained with fMRI connectivity (for identical subjects, subject measures and number of principal components entering CCA), see Figure 1S5A. Also, with regard to explaining the full set of subject measures, the MEG connectivity used here performs similarly well as the fMRI-based connectivity (Figure 1 S5B). Interestingly, what was also observed is that an entirely unresolved, i.e. one-dimensional broadband connectivity model performs less good in terms of maximum alignment and explained variance of subject measures. Furthermore, even when using the maximum spectral resolution available (20 2-Hz wide bins instead of 5 bands), the results were not outperforming the 5-band model.

## 3. Discussion

In summary, we have shown here that exploiting the spectral richness of MEG connectivity patterns by using a spectrally resolved connectome is useful to reveal a brain-behaviour mode not unlike previous results uncovered via hemodynamic connectivity metrics. Beyond simply reproducing, our results demonstrate the existence of functionally distinct networks across a broad range of conventional frequency bands. Even neighbouring bands, e.g. alpha and beta, can exhibit opposite roles with respect to a brain-behaviour mode arranging behavioural measures on a positive-negative axis.

Regarding the approach taken in this study, the concept of spectrally resolved connectivity, which can also be considered neuronal communication at multiple time scales or channels, is not new. Some models of how neuronal assemblies communicate have focused on activity confined to certain frequency bands^40^, while more recent perspectives also incorporate multi-channel or multi-frequency scenarios^15^. Furthermore, empirical studies, mostly MEG, have also increasingly shown connectivity and networks at multiple frequencies or showing features or states with specific spectral features or profiles^41,42,43,44^) and first reviews about the accumulated literature emerge ^45^. What was hitherto missing was a functional characterization of these spectrally resolved features or parameters, due to the absence of an attested link to behaviour. Here, to the best of our knowledge, for the first time, we have demonstrated the nature of the relationship between a spectrally resolved connectivity estimate to not just one, but an entire battery of behavioural and other subject variables.

Although the classification of brain activity into predefined frequency bands might be an oversimplification of the richness of neuronal dynamics in the brain, it can be a useful one. The fact that using these predefined, canonical, spectral ranges and the corresponding connectomes reveals a global brain-behaviour mode with little overlap between bands lends support to the idea of spectrally confined, functionally meaningful networks. This is also supported by our results, in a two-fold manner. First, the pure broadband connectivity (i.e. spectrally unresolved) approach yielded a brain-behaviour mode that was both less potent in explaining variance in behaviour than the spectrally resolved approach. On the other hand, the fully (i.e. 20-bin) spectrally resolved connectome also failed to reveal any additional insights into brain-behaviour associations compared to the approach conventionally assuming the existence of 5 bands. Of course, this does not mean that deploying the standard 5-band model is the optimum for identifying brain-behaviour links, but it lends support to the existence of well-defined and functionally distinct frequency ranges and the networks that emerge from them.

Naturally, the approach presented here can be extended in several ways. For example, our results do not rule out the existence nor the utility of more complex relationships in the time-frequency domain such as cross-frequency coupling or other types of neuronal communication that are less confined to spectral boundaries. Further studies looking into this will be needed to fully uncover the role of interactions in neural transient dynamics.

Further, the strong preference for either positive or negative relationships with the mode, corroborated by numerous control analyses, seems to be supportive of the existence of different ‘channels’ of neuronal communication, characteristic of and confined to these frequency bands and the edges involved. Furthermore, we do not find that a single frequency band completely dominates the CCA derived brain-behaviour mode. Rather, analyses of both the functionally relevant patterns of connectivity - the EMs - as well as the accumulated EM maps, point to a relatively distributed and balanced contribution of network components indicative of behaviour.

Another point of interest is the fundamental and qualitative difference between the observed EMs in alpha and beta-bands. While both rhythms, at rest, express functional connectivity network patterns that are relatively amenable to interpretation^32,41,44^ – they resemble visual or motor related networks - the resulting EMs, for the observed mode at least for beta, are much more frontally or prefrontally localized and can be less easily assigned directly to sensorimotor functions. In addition, higher beta-band connectivity is beneficial, while higher alpha connectivity (in the predominantly visual nodes implicated by CCA) is apparently deleterious for cognitive performance linked to the observed mode. This implies that higher connectivity strength is not always better, but its impact on the mode (and thus behaviour) appears to be a function of frequency. The finding of negative edge modulations in the alpha range may appear somewhat surprising, however, all additional control analyses we had performed support our finding of posterior, alpha-band specific connectivity was detrimental to cognitive performance. With regard to existing studies that focused on alpha-band connectivity, most of them were examining patient populations. Of these studies one study did – in line with our findings – report increased occipital alpha connectivity in schizophrenic patients compared to healthy controls^46^, which is interesting because schizophrenia often involves cognitive impairment. Other studies dealing with cognitive impairment arising from Alzheimers’ disease^47,48^ reported decreased alpha-band connectivity related to cognitive deficits, but more in frontal or unspecific regions. Thus, while there are some clinical studies that focused on alpha-band connectivity (or other bands), we are not aware of a healthy M/EEG cohort where a similar large battery of behavioural variables was collected. However, the study using HCP fMRI-derived connectivity found also most of their negative edge modulations in early visual areas being in line with our findings^5^. This, together with the observation that CCA identified a MEG-fMRI link of connectivity patterns (see Methods, “Validation”) lends further support to our findings.

The fundamental issue of ‘good’ and ‘bad’ brain patterns is also related to another line of research, which has investigated how variability predicts healthy ageing and cognitive performance^49,50^. While the approach employed here emphasised connectivity rather than variability per se, the two concepts are not entirely unrelated. For example, neuronal variability (or activity in the widest sense) is a necessary pre-condition for meaningful connectivity. What directly links our results to the concept of variability, is that the nodes implied in our functionally relevant networks (as seen in the accumulated EMs, for example) differ from others in their degree of envelope modulation at rest, which may be a sign of increased information transmission or communication. What is particularly noteworthy is that this level of modulation is associated with this node being important or not; it does not predict whether this variability is good or bad – this is rather dependent on the frequency band the network is operating at.

The apparent existence of multiple, functionally-relevant frequencies opens up the possibility of multiplexing, which is an elegant concept of how communication and information transfer may be realized in the human brain^34,51^. While we cannot completely exclude such a scenario, our results do not support this scenario. Rather, we obtained evidence for spectrally distinct networks, that are also spatially distinct in their network characteristics. The distinct character of these behaviourally relevant components may not necessarily require multiplexed communication. This highly complementary nature suggests that the idea of frequency bands is not just an entirely artificial construct, but seems to delineate different functional networks. It is important to note that the networks that are reflected by the identified edge modulations here are different from purely resting state networks that show coherent behaviour and as such are considered connected. The networks that are reported here are rather characterized by their similar functional meaning with respect to the identified brain-behaviour-mode. The patterns we find, i.e. the spectrally resolved edge modulations implicated in the canonical brain-behaviour mode, are for the most part spatially distinct patterns and do not show strong overlap across spectral boundaries.

In conclusion, we have provided evidence that electrophysiologically derived and spectrally resolved connectivity present in MEG resting state data can be used to index the ranking of individuals across a large range of subject measures. It exhibits highly structured patterns that are functionally relevant and provides sensitivity comparable to rs-fMRI derived networks in explaining variability of subject measures across a wide range of different features and domains. Furthermore, the additional dimension afforded by spectrally resolving connectivity measures opens up new avenues into a better and more holistic understanding of the roles and contributions of brain rhythms at rest that ultimately will help cast light on the intrinsic features of the brain that determine positive and negative cognitive and behavioural traits. Given recent breakthroughs in using frequency-tuned transcranial electrical stimulation techniques to enhance cognition^52^, outlining the roles of frequency-specific connectivity networks is all the more timely, and provides a basis for considering potential network-level targets for novel neuromodulatory interventions.

## 4. Methods

### Subjects and data

Resting state MEG data from the Human Connectome Project were used for this study. 89 subjects had complete resting state recordings (i.e. 3 recording sessions of about 6 mins). Mean age: 29 +/- 4 years, Sex distribution: 41 female / 48 male. Subjects were recorded in supine position and instructed to remain relaxed, with eyes open.

### Subject and behavioural measures

Following along the lines of a previous study showing CCA derived brain-behaviour mode with fMRI^5^, we extracted a large number of subject measures from the HCP data set. We excluded gender related subject measures and structural measures, but kept all other measures (cognitive, emotional, sensory performance tests, psychiatric and personality tests, family history of mental or other disorders, consumption of alcohol, tobacco or other drugs, in-scanner (fMRI) task performance). From those, we excluded variables that did not fulfil the following criteria:

1. Less than 80% of values in a subject measure should have same discrete values
2. At least 49 out of 89 subjects should have non-missing values for a given subject measure (compared to previous studies (REF), we chose this conservative number since our data set is considerably smaller).
3. The maximum value of any given value across subject measures should not exceed 100 times the mean of the group.

These exclusion criteria resulted in 131 behavioural variables that were used for further analysis.

We also deconfounded, i.e. regressed out, effects of age, handedness, gender, height, blood pressure (systolic and diastolic) and brain volume for all subjects in order to remove connectivity related effects mediated by these that might distort and confound connectivity measures (independent of the imaging modalitiy, i.e. MEG or fMRI, since both were used in the control analyses). Finally, all variables were further subjected to rank-based inverse normal transformation to ensure Gaussianity of distributions and demeaned. The general approach followed preprocessing of behavioural variables as previously conducted for an fMRI derived brain-behaviour mode ^5^.

### MEG pre-processing and parcellation, source leakage correction

MEG acquisition. Magnetoencephalography data was recorded by a whole-head MAGNES 3600 (4D NeuroImaging, San Diego, CA). Data were acquired in three runs of resting state recordings, lasting 6 minutes each. The MEG recorded from 248 magnetometer sensors, with 23 reference channels. Sampling rate was at 508.63 Hz and data were down-sampled for further processing to 200 Hz. The standard pre-processing pipeline as offered by the HCP (‘tmegpreproc’) was used which applied independent component analysis to remove potential artefacts from ocular, muscular or cardiac sources. For source reconstruction, single shell volume models were used, based on the individual anatomical MRI T1 images provided by the HCP consortium. Linearly constrained minimum variance (LCMV) beamforming was employed onto a regular 3D grid in normalized MNI source space with a resolution of 8mm^3^ using normalised lead fields and data covariance estimated in the 1-48Hz broadband frequency range. Source activity was normalised by the power of the projected sensor noise. Using PCA, the first principal component at each 3D source voxel location was extracted resulting in 5798 (1D) source-voxels. These were parcellated into 100 source parcels using a semi-data driven parcellation approach. Parcels were derived on the basis of a 246-region anatomical brain atlas^53^ that covers the two hemispheres (including subcortical structures). However, at this resolution, parcels are sometimes too closely related, resulting in rank deficiencies in the covariance matrix which in turn makes the following source leakage correction infeasible. Reducing them to 100 parcels averts this problem. In order to reduce the number of parcels optimally, we performed a k-means clustering of group-(and session) average correlation matrices of the raw time-series (parcellated into the initial 246 parcels per subject and session). This clustering identified the parcels that were correlated the most (in absolute terms) and merged them subsequently into a set of 100 parcels. Parcellation at higher dimensionality, e.g. with N=200, lead to failure of the multivariate source leakage correction needing full rank covariance matrices. A visualization is provided in Supplementary Figure 1, for a labelled list of parcels see Supplementary Table 1).

This parcellation is different to that used to analyse the relationships between fMRI connectivity maps and behaviour ^5^. We chose to use this data-driven customized parcellation based on the rationale that the fMRI-derived parcellation is a spatial ICA capitalizing on the large amount of spatial information available in fMRI data, sometimes resulting in spatially fine-grained components. However, since this level of spatial resolution is not always given in MEG data, thus, similar to the higher-dimensional BrainNetome^53^ derived parcellations from above (with N>200), rank deficiency of the parcel-time series can render the source leakage correction infeasible. For the control analyses that employed fMRI-derived measures of functional connectivity we employed the previously published approach^5^, but chose the lower resolution of N=100, corresponding to the dimensionality chosen for MEG analysis.

To obtain parcel-time series, the first principal component over all voxels belonging to any given parcel was extracted. After extraction, resulting parcel time-series underwent reduction of source leakage by using a multivariate leakage reduction approach^54^, effectively orthogonalising all parcel time-series and removing zero-lag correlations.

### Analysis of functional connectivity

After source-leakage correction, we estimated functional connectivity (FC), separately for each individual and session. Functional connectivity was estimated using the OSL toolbox (Oxford Software Library), freely available at https://github.com/OHBA-analysis/osl-core. We first computed envelopes in 20 contiguous frequency bins, with a frequency range of 0.5 to 40 Hz, and a resolution of 2 Hz, using the resulting envelopes in each bin after Hilbert transformation of the narrow-band filtered time-series. Within each bin, full linear Pearson correlation was computed between all possible pairings of parcel-based envelope time-series, resulting in a 100 × 100 connectivity matrix for each resting state run, frequency bin and subject. After FC estimation, all three runs were then averaged to increase stability of FC estimations. After averaging, connectivity measures, i.e. the correlation coefficients were first Fisher z-transformed.

We observed that functional connectivity in spectrally resolved MEG data is not always independent of spectral power or frequency, but that higher spectral power is linked to higher group-average envelope correlations, with connectivity being stronger (on average) in lower frequency bands (which tend to have higher power due to the well known 1/f behaviour of human brain power spectra). In order to avoid that our findings were not simply confounded by these effects, we removed this systematic variation by pair-wise regressing out of spectral power from connectivity measures (i.e. envelope correlation coefficients) from all possible pairings (i.e. edges). This resulted in subject-specific connectivity estimates without power-induced low-frequency bias. After this, we removed the other previously defined confounds (see Methods, “Subjects and Behavioural measures”). Subsequently, the resulting connectivity matrices were renormalized to ensure zero mean and unit variance.

For further analyses we chose to further reduce the number of frequency bins to 5 bands, so we averaged the initial spectrally resolved connectivity data to approximately follow conventional frequency band arrangement: delta-band [0.5 - 3 Hz], theta-band [3-7 Hz], alpha-band [7-13 Hz], beta-band [13-25 Hz] and lower gamma-band [25-40 Hz]. Before further analysis, we extracted the upper triangle from each symmetric connectivity matrix, concatenated the resulting matrices (5×4950×89) frequency bands (yielding a 24750 × 89 matrix FC1).

### Dimension reduction and analysis of brain-behaviour mode by means of canonical correlation analysis (CCA)

Behavioural as well as connectivity data were subjected to principal component analysis (PCA). For connectivity, PCA is performed on the concatenated connectivity data sets (i.e. on the 24750 × 89 concatenated connectivity matrix (FC1), see Figure 1 S4 for visualised approach), for behaviour it is performed on the 131 × 89 behavioural data set. For both data sets, we retained 22 components per subject. We chose this specific dimensionality because it is low in comparison to the sample size (22 PCs vs 89 subjects, reducing the chance of overfitting (see also Methods “ alidation” section for more details) in downstream analyses) and still ensures sufficient detail retained in the reduced model. After reduction to 22 components, 74.1% and 64.5% of the variance was explained for the behavioural data and the connectivity data, respectively. This resulted in a 22 × 89 PCA-reduced connectivity matrix, FC2 and a 22 × 89 PCA-reduced behavioural matrix (B2).

After PCA-derived dimension reduction we applied canonical correlation analysis (CCA) to the PCA-reduced data. CCA is a method that identifies the linear combination in each of two sets of features that transforms them to produce a maximal match of the two sets. The linear combinations that maximize correlation between the two sets of features or matrices are typically referred to as “modes”. We employ CCA here to examine whether there is such a match, or mode, of brain-behaviour population covariation for MEG functional connectivity and behavioural data. In contrast to fMRI approaches, MEG contains the additional dimension of frequency, here characterised in five frequency bands CCA is then performed on these two identically-sized matrices (FC2and B2), finding the linear combination of weights (or mode) that transforms each of the two matrices into new matrices (FC3 and B3) where each row is maximally similar to each other. Each row in FC3 and B3 corresponds to a newly formed canonical variate that is orthogonal to all other canonical variates within FC3 and B3, representing the linear weighting of connectivity (FC2) and behavioural features (B2) to best align brain and behavioural covariation across subjects. Each row with their corresponding canonical variates corresponds to a “mode”, with decreasing degree of correlation. Consequently, CCA result in a maximum of 22 canonical modes with 2×22 canonical variates that are also orthogonal within FC3 and B3. ach mode’s statistical significance, that is, the significance of the correlation between rows in FC3 and B3 is then established by permutation testing (with n = 10000), which corrects for multiple comparisons. For permutation testing, subject labels were swapped while respecting family structure in the data. For running CCA and permutation testing we used an adapted version of the scripts made available in a previous CCA study^5^.

### Visualisation of post-CCA results

For characterization of the observed CCA mode(s) with respect to the underlying behavioural variables and connectivity, the resulting behavioural CCA weights (i.e. one per subject, n=89 in total) were correlated with the full, deconfounded behavioural variables (i.e. 131 × 89), and the mode-derived connectivity weights (n=89), in turn, were correlated with the full, deconfounded individual FC networks (i.e. the original 5 × 4950 x 89 connectivity matrix). For visualization of connectivity and the resulting edge modulations, we used the freely available Circos software suite^55^ (http://circos.ca/software/download/circos/). We also derived accumulated EMs from the data where one dimension of the 100×100 EM matrices was collapsed by averaging over connections. For visualizing the results, we used the parcellation functionality of OSL (Oxford Software Library), available at https://github.com/OHBA-analysis/osl-core.

### Validation of CCA results

CCA is highly efficient in maximizing correlation between data sets, finding the features (or a linear combination thereof) that are maximally aligned. However, this naturally leads to concerns of overfitting. While it already has been shown that similar approaches have yielded robust and stable results for larger data sets (with a similar parametrization of CCA input regarding the ratio of features to sample size), the current dataset is smaller. Thus, while we are confident that our choices, e.g. a low number of retained components, a relatively modest number of parcels etc. are conservative and ensure robustness, we performed several additional control analyses to rule out overfitting and to add confidence in the stability of the presented results. The following analyses were performed:

1) In a first cross-validation test, we split each subject’s MEG resting state data set into two parts: One part consisted of the first two resting state sessions, the other part consisted of the last session. CCA was applied to the first, acting as a training set - identifying the best matching mode in the first two sessions - whereas the second data set (the remaining resting state session) acted as test set - by applying the identified weights from the training set on the test set, effectively predicting subject and connectome weights for the test set. These predicted subject measure and connectome variates for the test subjects were then correlated with each other to obtain an estimate of how well the weights identified by the CCA on the training set still work in the test set. Mean correlation in the first mode, i.e. between the first canonical variate of subject measures and the corresponding first canonical variate of connectome data in the training set (FC3_(1)train_ vs B3_(1)train_) was 0.9X, while the brain-behaviour correlation coefficient in the test set (FC3_(1)test_ vs B3_(1)test_) was at 0.79. As a reference, the mean correlation of the permutation null distribution in the test data set was 0.02, with a standard deviation of 0.11. At the observed correlation of r=0.79, significance of the observed correlation was at max p = 1×10^−4^ (the threshold for r at a 5% alpha-level (corrected) being r = 0.21).
2) For a second, complementary cross-validation test, we used a leave-one-subject-out approach to test whether there is overfitting on a cross-subject level. Accordingly, CCA was applied to a first training set of all subjects apart from the left-out subject (n = 88) - identifying the best matching mode in these subjects - and applying these weights (that form the first canonical variates) on the left-out subject set effectively obtaining predicted subject and connectome canonical variates for the left-out, test subject. Subsequently, this approach was repeated for all 89 subjects, resulting predicted subject measure and connectome scores for all subjects. These predicted subject measure and connectome scores for the test subjects were then correlated and tested for significance. For this type of cross-validation, correlation coefficient r in the first mode, i.e. between the first canonical variate of subject measures and the corresponding first canonical variate of connectome data in the training set (FC3_(1)train_ vs B3_(1)train_) was at r = 0.67. Mean correlation coefficient r of the permutation null distribution was <0.01, with a standard deviation of 0.11. Significance of the correlation of the first mode was at maximum of p = 0.001 (with a 5% threshold of a correlation coefficient r being 0.1930).
3) In this control analysis, we tested the stability of our spectrally resolved approach with regard to the resulting band-specific edge modulations, i.e. the functionally relevant networks identified in each of the five bands. To do so, we now split the connectivity data, for each subject, into 5 sets, each containing only a single frequency-band specific connectivity matrix and then ran the described PCA-CCA pipeline separately on those. We then compared the resulting separate edge modulation maps (EM_sep_) with the corresponding band-specific edge modulation maps of the original, integral CCA analysis (EM_orig_), where all 5 bands entered CCA in their entirety (we chose this correlation as a test criterion here, since in this case neither canonical subject measure variates nor functional connectivity variates (FC3, B3) are directly comparable to the one integral main analysis. The resulting correlation coefficients between EM_sep_ and EM_orig_ were: in delta band 0.76, theta band 0.48, alpha band 0.80, beta band 0.48, and in the gamma band 0.32. For interpretation of these results it needs to be said that a holistic, integral CCA of all five band-specific connectomes is not expected to give identical results in each band as the band-separate analysis in the control condition, but it answers the question whether the general relevance, for example the quality of each identified band-specific network is preserved (see ‘good or bad’ section in Discussion). All bands show, on average, the same qualitative relevance – being predominantly either good or bad, as in the original analysis. For an exemplary illustration of the alpha-band edge modulations in analysis 2 and 3 see Figure 2 S1.

For the remaining control analyses we will present the validation and control analysis results in the following order: CC1, CC2 and CC3; with CC1 being the correlation coefficient between the first canonical variates, representing the first canonical mode in FC3_orig_ and FC3_control_, and CC2 being the correlation between first canonical variates in B3_orig_ and B3_control_, while CC3 is the resulting brain-behaviour correlation (FC3_alternative_ vs B3_alternative_) in the first canonical mode:

4) In this control analysis, we ran a CCA on the same 89 subjects, where we only used fMRI derived connectivity measures, analogous to previous reported approaches. Similar to our main result and previously reported fMRI results, we found one significant mode arraying subjects on a positive-negative axis of behavioural variables. Resulting connectome (FC3_MEG(1)_) and subject measure scores (B3_MEG(1)_) for the first canonical mode of the MEG derived main analysis (5-bands), and the fMRI derived canonical scores (FC3_FMRI(1)_ and B3_FMRI(1)_ for the first mode correlated by CC1 = 0.26 and CC2 = 0.27, respectively. When comparing these fMRI derived CCA results against the full 20-frequency-bin MEG-CCA correlations were slightly higher: CC1 = 0.3 and CC2 = 0.32, respectively. While these correlations seem relatively low, qualitatively, the fMRI-derived CCA approach identified a positive-negative axis within the first canonical mode similarly to previously reported and identified here by the main MEG CCA analysis, see Figure 1 S3 for the behavioural variables associated with and their ranking within this mode. The resulting first canonical mode yielded a correlation coefficient of CC3 = 0.92.
5) We compared the CCA outcome of our 5-band approach to the outcome using the initially estimated, ‘fully’ spectrally resolved 20-bin connectivity data (see Methods above). Correlations of subject measure scores and connectome scores (i.e. FC_orig_ vs FC3_control_ and B3_orig_ vs B3_control_) were at CC1 = 0.8 and CC2 = 0.85, respectively. The resulting mode gave a brain-behavior correlation coefficient of CC3 = 0.93.
6) In this control analysis we did not perform source leakage. Correlations between the original, full-source leakage corrected model and the non-corrected model were: CC1 = 0.25 and CC2 = 0.27 for the first canonical variates (FC3_orig_ vs FC3_control_ and B3_orig_ vs B3_control_). Resulting brain-behaviour correlation was CC3 = 0.89.
7) A model where we only used 50 parcels (fMRI-derived parcellations from the HCP initiative, HCP1200 release available from https://db.humanconnectome.org/data/projects/HCP_1200) for parcellation of MEG data instead of the 100 in our original analysis, correlation to the original canonical variates were CC1 = 0.14 and CC2 = 0.16 respectively (FC3_orig_ vs FC3_control_, B3_orig_ vs B3_control_). Resulting brain-behaviour correlation was CC3 = 0.90.
8) In another control analysis, we chose an approach where connectivity was defined as broadband connectivity (1-40Hz) only, so instead of several bands or bins there was only one connectivity measure per parcel. Correlations with the original canonical variates (FC3_orig_ vs FC3_control_ and B3_orig_ vs B3_control_) were CC1 = 0.36 and CC2 = 0.38. Resulting brain-behaviour correlation was CC3 = 0.90.

The correlation coefficient of each identified first mode for control analyses 4-8 can be found visualized in Figure 1 S5A.

9) In a last control analysis, we tested whether MEG and fMRI connectivity show some overlap in explaining variance across the subject population, i.e. we tested the link between these two independently acquired and modality-specific connectomes. This test was performed by using only connectivity data as input to the CCA, i.e. one set of variables being the fMRI connectivity data while the other one was the MEG based connectivity data (from the same subjects), asking for the existence of a connectome-connectome mode of population covariation. As result, we observed one significant mode, tying together MEG and fMRI connectivity across subjects was identified (resulting brain-behaviour correlation coefficient in first mode was r = 0.92). Apart from the demonstration of the existence of such a link, we did not perform further analyses to look into spatial or spectral components of this link or visualize them, since it was beyond the scope of this study.

We also computed the explained variance of the first and all following canonical variates, i.e. the first and significant CCA mode until the last computed mode with regard to the set of behavioural variables. Results are shown in Figure 1S2, demonstrating that the first canonical mode explains significantly more variance in behaviour than the permutation based null model where we swapped subject label n=1000 times. For a comparison of the explained variance in the control analyses 4-8, see also Figure 1S5B.

## 5. Acknowledgments

Funded by the Swiss National Science Foundation (grant: PP_163726 awarded to A.H-A). Data were provided by the Human Connectome Project, WU-Minn Consortium (Principal Investigators: David Van Essen and Kamil Ugurbil; 1U54MH091657) funded by the 16 NIH Institutes and Centers that support the NIH Blueprint for Neuroscience Research; and by the McDonnell Center for Systems Neuroscience at Washington University. We are grateful to Steve Smith and colleagues for sharing the code of their HCP-fMRI CCA study^5^ online.

## 6. Data availability

The sensor space MEG resting state data and the corresponding subject measures are available online on https://db.humanconnectome.org, however they are not publicly accessible without registration. Due to privacy concerns, access to these data needs registration and approval by the HCP consortium. All authors have been approved and have accepted the terms of use for the open and restricted part of the HCP data.

The processed and derived data (functional connectivity, CCA results etc.) that support the findings of the study can be made available upon reasonable request to the corresponding author (R.B.). Since derived from the HCP data, the processed data are not publicly available due to them containing information that could potentially comprise research participant privacy and / or violate HCP terms of use and thus will be only shared accordingly.

## 7. Code availability

The code used in this study uses several publicly available toolboxes and software. A previous implementation of CCA with respect to HCP functional connectivity and subject measures has been used elsewhere^5^ and can be found on https://www.fmrib.ox.ac.uk/datasets/HCP-CCA/. Circos software is available on http://circos.ca/software/download/circos/. OSL software is available on https://github.com/OHBA-analysis/osl-core/. Customized Matlab scripts for source reconstruction, use of PCA / CCA, post-CCA visualizations using OSL and Circos can be made available on request to the corresponding author (R.B.).

## Supplementary Figures

**Figure 1 S1.**
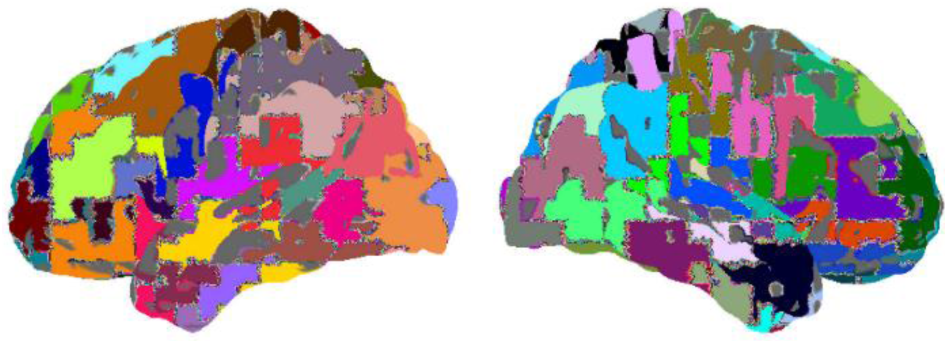
Visualisation of the used parcellation (n = 100, lateral view). Parcel identities can be found in Supplementary Table 1, with anatomical labels.

**Figure 1 S2.**
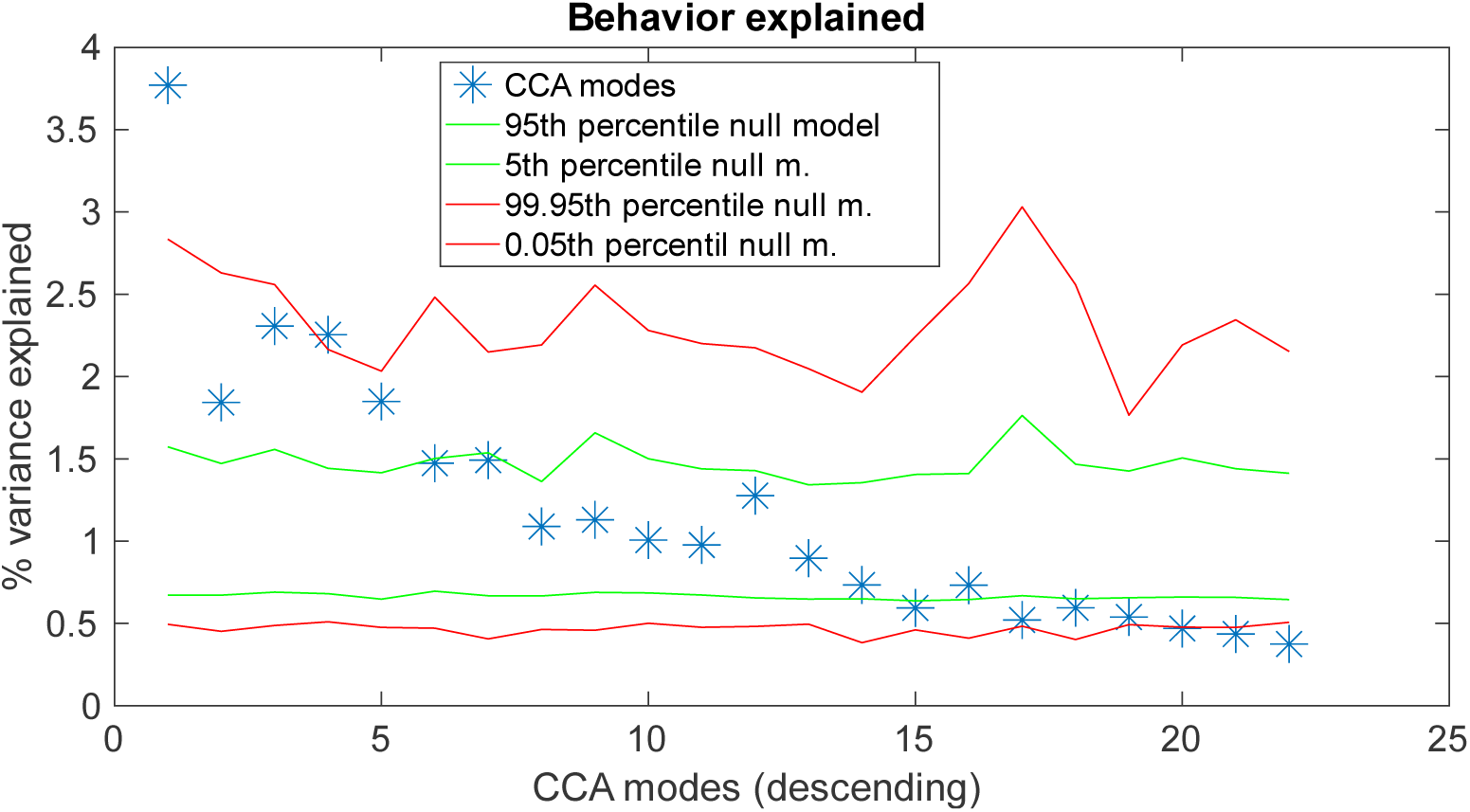
The amount of variance (%) explained by the first canonical variate (i.e. first mode) in the full set of behavioral variables. Permutation test n =1000. Green and red lines indicate differently thresholded top and bottom percentiles of permutation based null model.

**Figure 1 S3.**
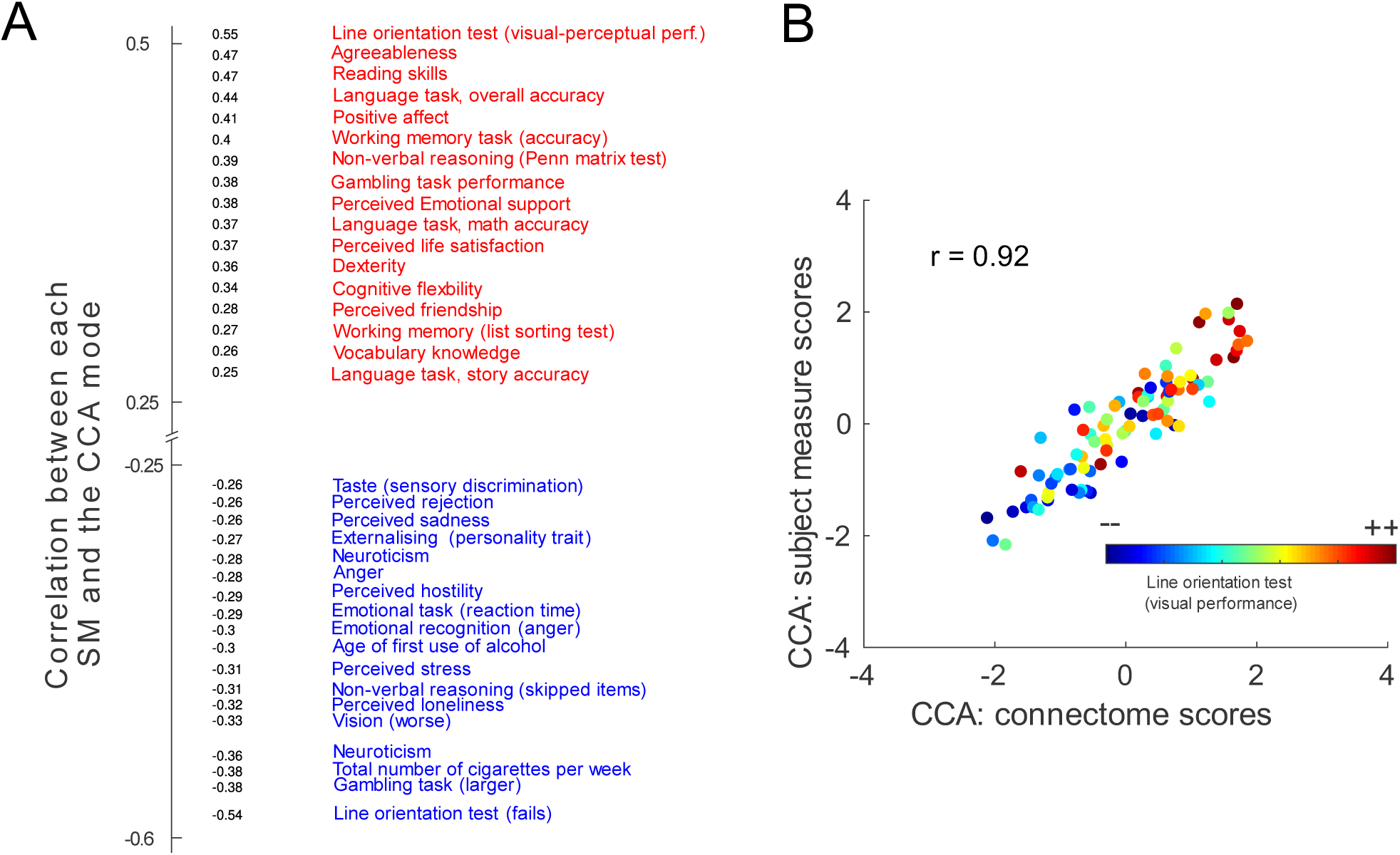
First CCA mode using fMRI derived connectivity (n=89). **A**. Behavioural ranking (thresholded at |rho|>0.25) and brain-behaviour mode visualization across subjects. **B**. Correlation of CCA-derived subject measure scores and connectivity scores for the first canonical variates identified, i.e. the first canonical mode. In colour, the behavioural score for the top-ranking behavioral variable in this mode, here line orientation discrimination, is shown per subject.

**Figure 1 S4.**
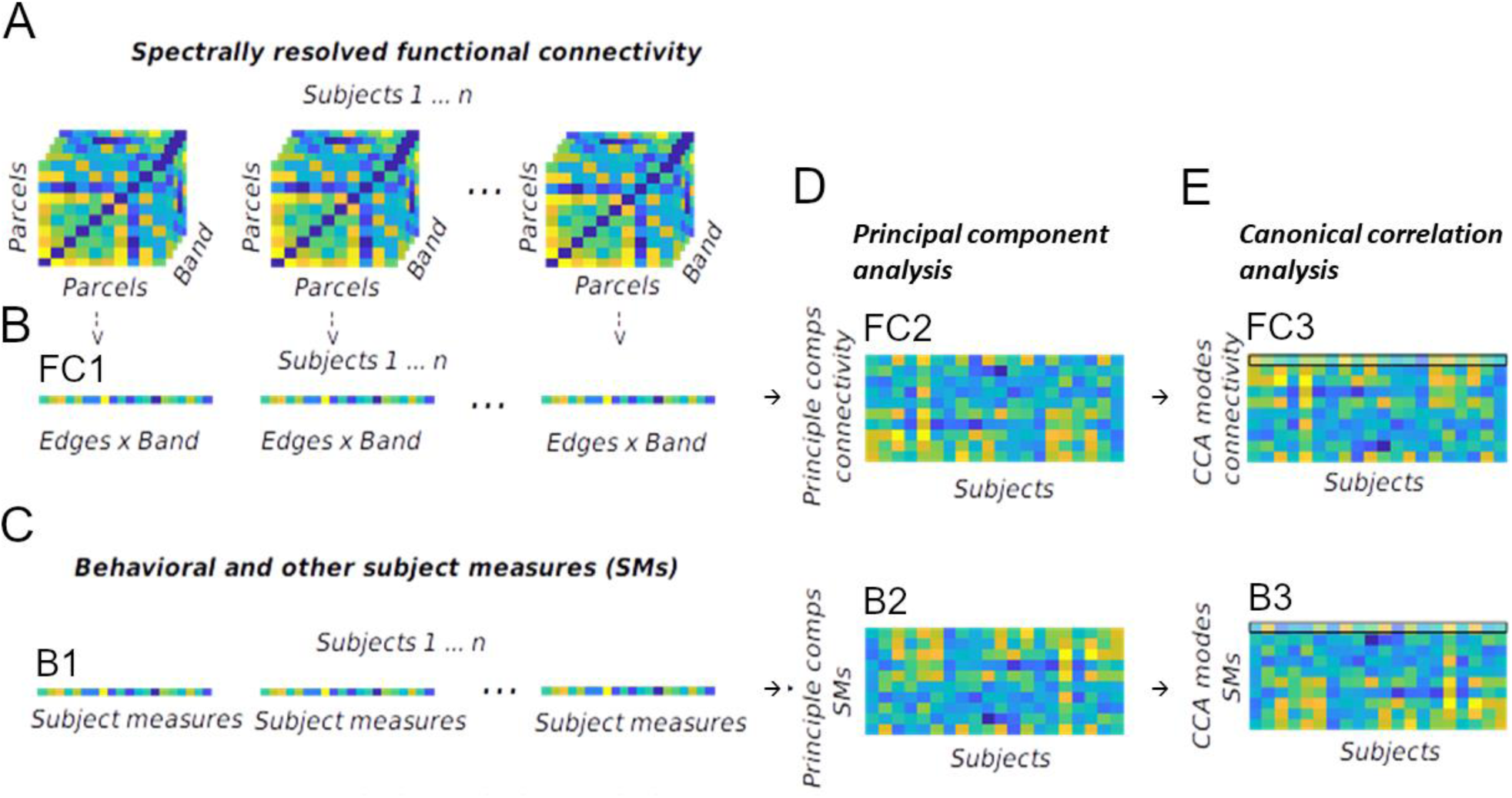
Illustration of analysis approach. Functional connectivity as defined by envelope correlations within five conventional frequency bands are extracted from n=89 subjects. For each subject, these spectrally resolved connectivity features are first concatenated (A, forming matrix FC1) and subjected to group PCA, retaining the first 22 principal components per subject (resulting in matrix FC2). The same approach is used for reducing dimensionality of the subject measures of all subjects (concatenated vector B1), retaining the first 22 principal components (resulting matrix B2). These two matrices (FC2 and B2) are then subject to canonical correlation analysis (CCA), which identifies within each matrix the optimal linear weighting to maximize correlation of features between the two sets of variables (i.e. brain vs behaviour), transforming matrices FC2 and B2 into FC3 and B3. There, each row represents one mode where the newly formed (i.e. linearly recombined) connectivity and behavior canonical variates correlate most strongly (first mode here is indicated by thin black line).

**Figure 1 S5.**
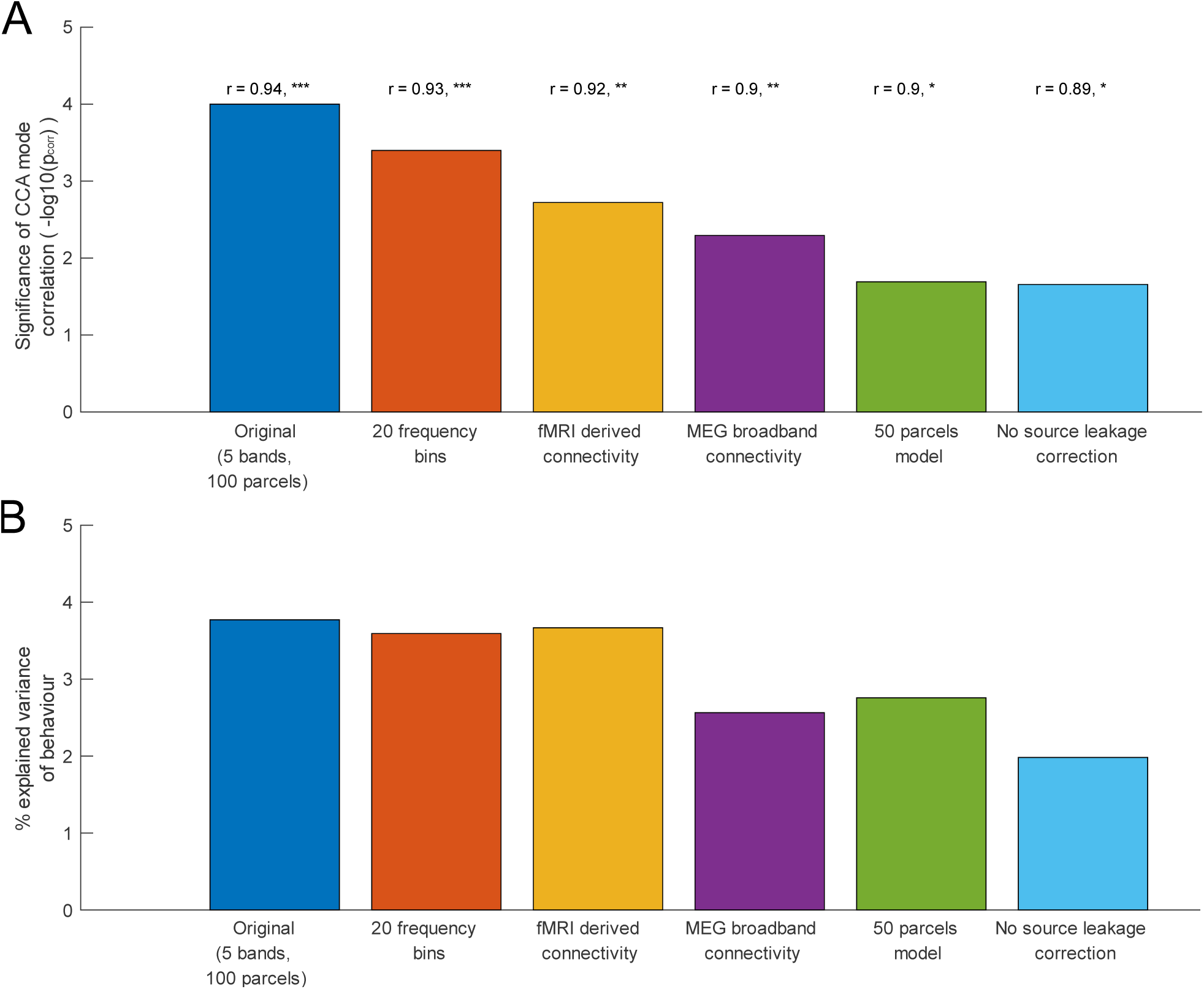
Comparison of different CCA analyses. **A**. Correlation of first canonical variates (first mode) of connectivity vs behaviour (y-axis show level of significance of correlation in negative logarithmic scale). **B**. Explained variance (in percent) of the behavioural variables in the first mode, same connectivity models as in A. In the order of appearance: 1 - Original analysis (5 frequency bands, 100 parcels). 2 - 20-bin resolved MEG connectivity data. 3 – fMRI connectivity used instead of MEG connectivity. 4 – Spectrally unresolved MEG broadband connectivity (1-40Hz) used, 5 - 50 parcel MEG connectivity data used. 6 – As original analysis, without source leakage correction performed. See also methods (“Validation analyses”, analyses numbered 4 to 8).

**Figure 2 S1.**
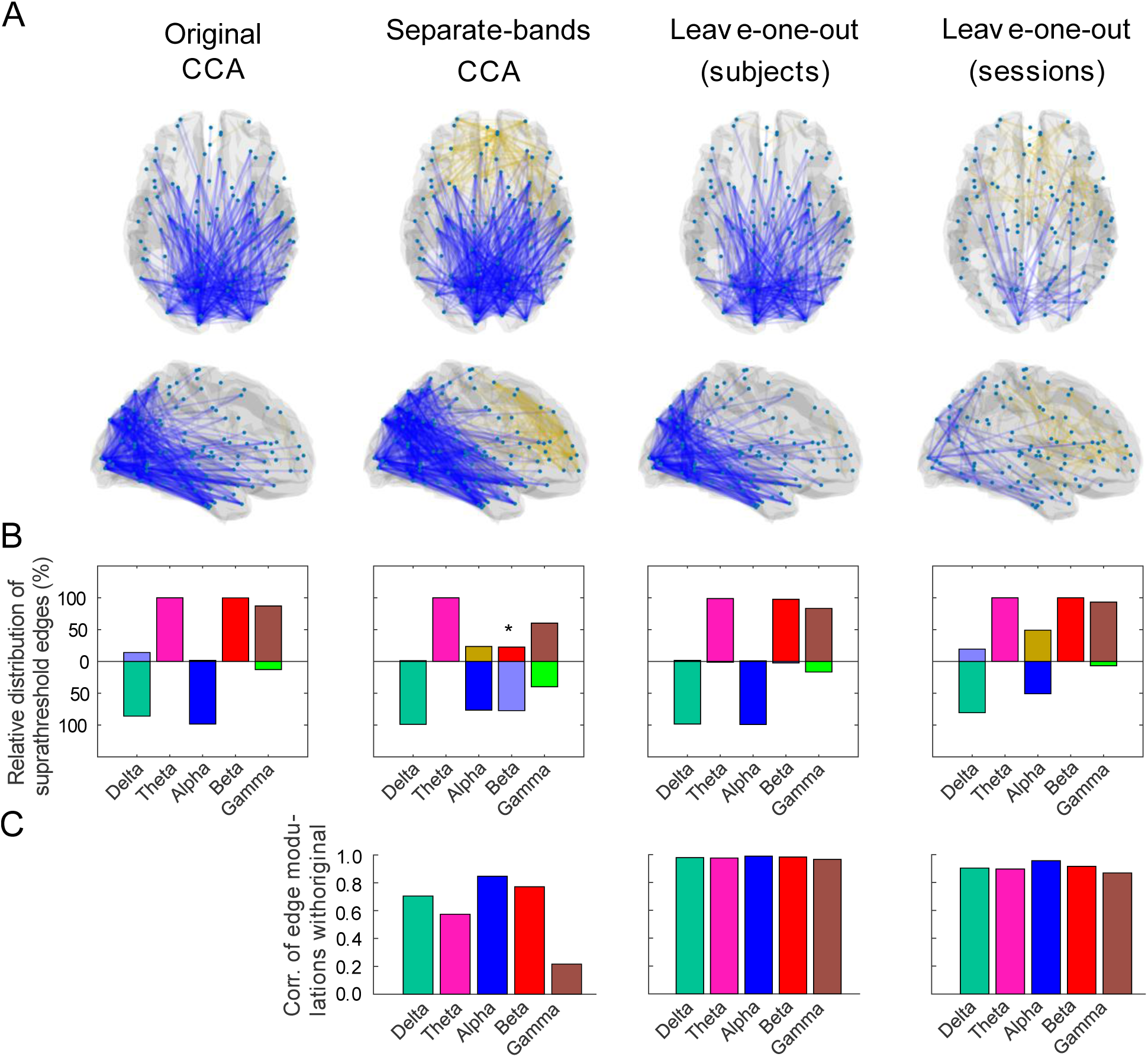
Stability of edge modulation results for different analyses **A**. Examplarily for the alpha-band, edge modulations for the original (main) analysis and three control analyses. First panel: Results from original analysis, visualizing the alpha-band edge modulations. Second panel: CCA has now been performed separately for each frequency band, here the edge modulation result for the alpha-band model is shown. Third panel: For this analysis, data were split into a training and test set, where the training was carried out on the first two resting state sessions and testing was carried out on the third session. Alpha-band edge modulation are shown for the CCA mode that was learned in the training set and applied to the test set. Fourth panel: In this analysis, data were split into training and test set by a leave-one-subject-out approach. Alpha band edge modulations are shown for the CCA mode as established for the left-out subjects (and their predicted canonical variates, see Methods for details). All alpha-band edge modulations show comparable patterns with the majority of supra-threshold patterns being negative, posterior edge modulations related to the identified CCA mode. **B**. Summary of the distribution of suprathreshold edge modulations across all frequency bands. Complementary to first row, this shows all frequency bands from delta to gamma. Almost all analyses behave the same way, apart from the beta-band in the frequency-band separate CCA analyses (where beta band CCA did not yield any significant mode). **C**. Similarity of edge modulation patterns to the original main analyses (first panel, Figure 1 and Figure 2). Here, correlation was performed over the whole 4950-element EM vector in each frequency band. High correlations indicate good correspondence with original EM patterns from the main analysis (as visualized in Figure 2). See also methods (“Validation of CCA approach”, analyses 1-3).

**Supplementary Table 1.**
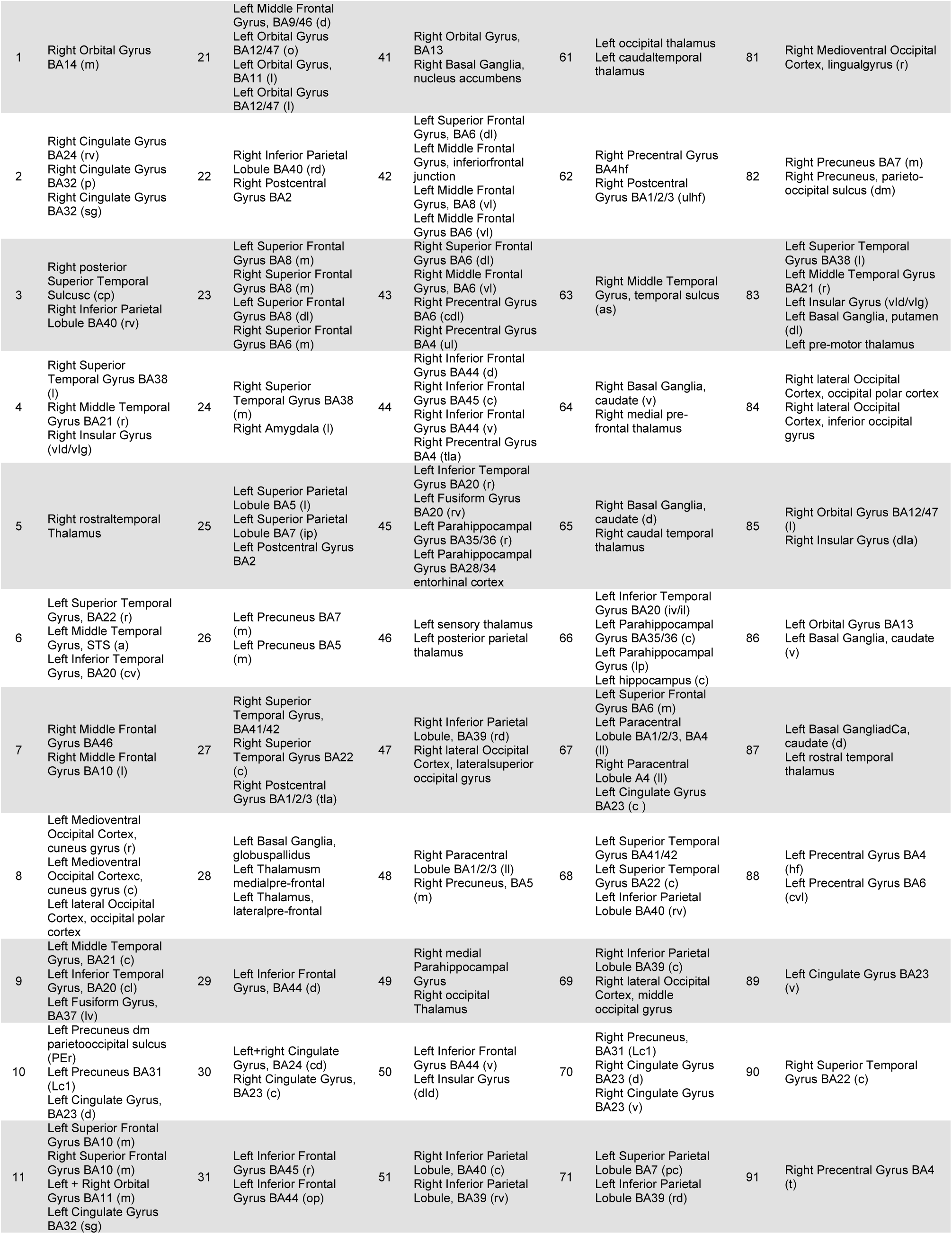
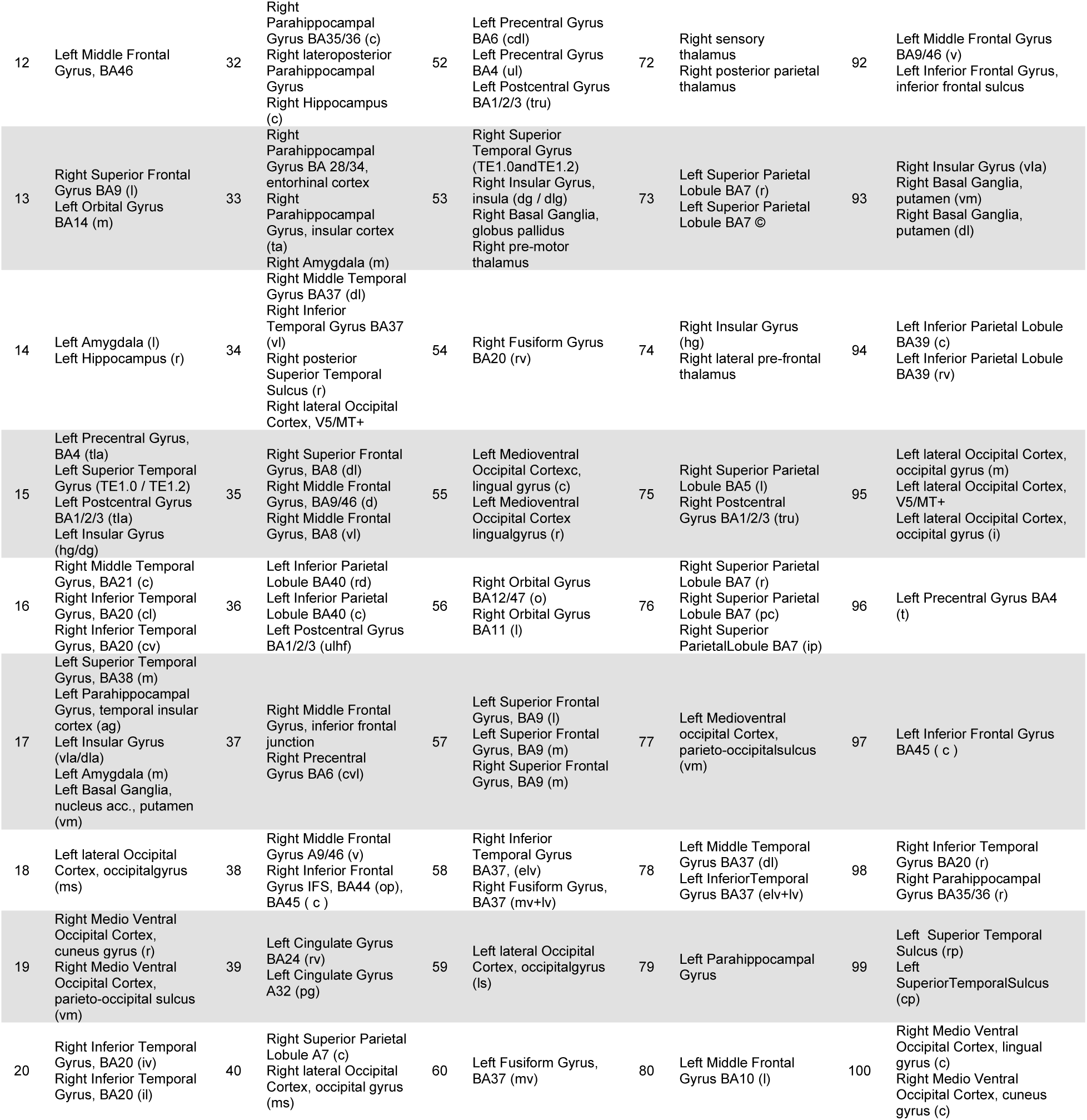
Table showing the parcel identities with anatomical labeling derived from the BNA atlas^53^. Mention of several areas means the parcel has been created by merging these originally separate BNA parcels into a single parcel. Abbreviations: m = medial, l = lateral, r = rostral, a = anterior, p = posterior, v = ventral, c = caudal, i = inferior, o = orbital, p = pregenual, ag = agranular, rv = rostroventral, sg = subgenual, cv = caudoventral, cl = caudolateral, dg = dorsal granular, lv = lateroventral, dm = dorsomedial, vm = ventromedial, tla = tongue and larynx area, ms = medial superior, iv = intermediate ventral, rd = rostrodorsal, cd = caudodorsal, rp = rostroposterior, op = opercular, vla/d = ventral a/dysgranular, da = dorsal agranular, ms = medial superior, ulhf = upper limb, head and face region, op = opercular, pg= pregenual, ms = medial superior, ll = lower limb region, dld = dorsal dysgranular, iv = intermediate ventral, pc = postcentral, ip = intraparietal, vld = ventraldysgranular, vlg = ventral granular, dla = dorsal agranular, cvl = caudal ventral lateral, ta = temporal agranular, lp = lateral posterior.

